# Introducing flocker: an R package for flexible occupancy modeling via brms and Stan

**DOI:** 10.1101/2023.10.26.564080

**Authors:** Jacob B. Socolar, Simon C. Mills

## Abstract

1. Occupancy models are a widespread tool for analyzing biological survey data, but packages for fitting these models offer a limited variety of effects structures.
2. We developed `R`package `flocker` to connect occupancy models (single- and multi-species; single- and multi-season) to the uniquely powerful formula syntax of `R`package `brms`.
3. Using familiar formula-based syntax, `flocker` models can readily incorporate a wide array of effects structures, including phylogenetic random effects, splines, Gaussian processes, autoregressive structures, monotonic effects, and nonlinear predictors. These are available for use in formulas for occupancy, detection, colonization, extinction, and autologistic terms (as applicable to the model type). `flocker` additionally provides functionality for data simulation, posterior prediction, and model comparison, following well-documented statistical decisions that we put forward as best practices.
4. We anticipate that `flocker` will facilitate the work of practitioners who seek added realism in occupancy models. We further hope that `flocker`’s synthesis of these models will help inform best practices around occupancy modeling.

## 1 Introduction

Occupancy models are an essential part of the modern toolkit for analyzing biological survey data, enabling ecologists to estimate species distributions, habitat suitability, and population trends while accounting for imperfect detection (*1, 2*). At their core, occupancy models can be thought of as a pair of linked regressions with binary outcomes, the first predicting whether a species occupies a site, and the second predicting the detection history over multiple sampling occasions, conditional on the site being occupied. By assuming that occupancy does not change over repeat visits, these models disentangle occupancy versus detection probabilities (*3*).

Several software packages are now available to fit occupancy models via formula-based syntax (*4* –*8*). However, the flexibility of existing packages to specify rich effects structures remains limited, constraining researchers’ ability to capture the complexities of ecological data. To meet this need, we developed `flocker`, an R package that connects the powerful formula syntax of `brms` (*9, 10*) to occupancy model likelihoods.

On the backend, `flocker` fits models in Stan (*11*) via `brms`, a state-of-the-art `R` package for specifying Bayesian statistical models via formula-based syntax (*9*). Among similar packages, `brms` is unique for the sheer variety of available effects structures, including flexible correlation structures, phylogenetic random effects, splines, Gaussian processes, autoregressive structures, monotonic effects, nonlinear predictors, measurement error in covariates, etc (*10, 12*).

Fitting occupancy models via `brms` is achieved by passing custom likelihoods and data structures to `brms`, but the process is complicated by the fact that most classes of occupancy model do not to factorize at the row level (Box 1), necessitating bespoke data structures and custom post-processing functionality. `flocker` solves these problems by providing:

1. A data format that encodes the data’s inherent visits-within-sites structure in a dataframe that `brms` will accept as input (see [vignette here](https://jsocolar.github.io/flocker/articles/flocker_format.html)).
2. A formatting function that accepts data inputs in intuitive formats and reformats them to the `brms` compatible specification.
3. `brms` custom families that include Stan code capable of decoding this information and computing the likelihood.
4. Simulation-based calibration (*13, 14*) of custom families to confirm algorithms function as intended (see [vignette here](https://jsocolar.github.io/flocker/articles/flocker_format.html)).
5. Bespoke post-processing functionality to handle the visits-within-sites data structure for the purposes of posterior prediction, model comparison, and so forth.

`flocker` implements single-season models (*3*), multi-season models of several kinds (*15, 16*), and models with data augmentation for never-observed species (*17*), bringing the full variety of `brms` effects structures to the relevant distributional terms in each class (i.e. occupancy, detection, colonization, extinction, and/or autologistic terms). We encode the data structure via auxiliary columns in the dataframe passed to `brms`. Stan code to decode this information depends on the number of repeat visits (and in multi-season models on the number of seasons), and is generated on-the-fly at runtime. In this paper, we review concepts and introduce new terminology around occupancy models in Box 1. We then provide a minimal introduction to model-fitting in `flocker`, full descriptions of the available model types, and an overview of `flocker`’s post-processing machinery. Throughout, we discuss subtleties and points of confusion that arose during our work. The package is available on GitHub at github.com/jsocolar/flocker. Example code for all model classes and post-processing features is provided in the [tutorial vignette](https://jsocolar.github.io/flocker/articles/flocker_tutorial.html), while simulation-based calibration results for all model types are provided in the [SBC vignette](https://jsocolar.github.io/flocker/articles/sbc.html).

### Box 1

**Core concepts and terminology**

#### 1.1 Closure-units

All occupancy models assume that there is some level at which occupancy does not vary, with any variation in observed data attributed to the detection process. The details of this *closure assumption* depend on the type of occupancy model and the biological system that it represents. Single-season single-species models (including models augmented with never-observed species) assume closure within sites. Single-season multi-species models assume closure at combinations of species *×* site. Multi-season models assume closure at site *×* year combinations (species *×* site *×* year combinations in multi-species versions). To refer in a general way to the units over which closure is assumed, we use the term *closure-unit*.

#### 1.2 Likelihood factorization

Many statistical models have likelihoods expressible as the product of multiplicative terms, each representing one data point with its associated covariates, commonly represented as one row in the input data. We say that such models *factorize at the row level*. Standard single-season occupancy models factorize at the row level only when detection probabilities are modeled as constant across repeat visits within each closure-unit. When detection probabilities vary across repeat visits within a closure-unit (e.g. according to time of day or observer identity), multiple rows of detection covariates are required to compute the likelihood for a closure-unit, and the model does not factorize at the row level. In multiseason models, the likelihood factorizes at the level of a site (or a species *×* site in multi-species models), but not at the level of a closure-unit. In data-augmented multi-species models, the likelihood factorizes at the species level.

#### 1.3 Event covariates and unit covariates

Covariates that vary across replicate visits within a closure-unit cannot be used to model occupancy. We refer to such covariates as *event covariates*, as contrasted with *unit covariates* that are constant within closure-units. Both unit covariates and event covariates can be used to predict detection, but only unit covariates may be used to predict occupancy, colonization, and extinction. We refer to a model with no event covariates as a *rep-constant* model, as opposed to a *rep-varying* model with event covariates.

## 2 A basic example

Flocker can be installed and loaded via:

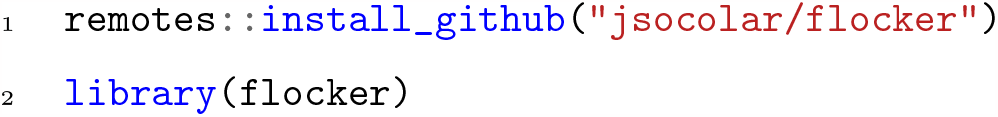

To fit an occupancy model, we first create a dataset and format it for use with `flocker`. General purpose data simulation is provided via `simulate_flocker_data()`, which by default simulates 30 species sampled at 50 sites using four replicate surveys (i.e. a single-season multi-species dataset). Non-default arguments simulate data from other likelihoods, including multi-season and data-augmented models. To pass data to `flocker`, we first pass the output from `simulate_flocker_data()` to `make_flocker_data()`, which repackages the data to the `flocker_data` format (see vignette here):

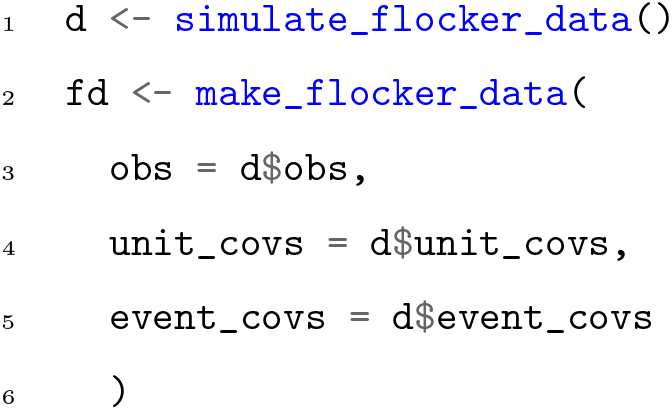

To fit a model, in this case a single-season multi-species occupancy model, we use the function `flock()`, which is capable of fitting all flavors of occupancy model described below.

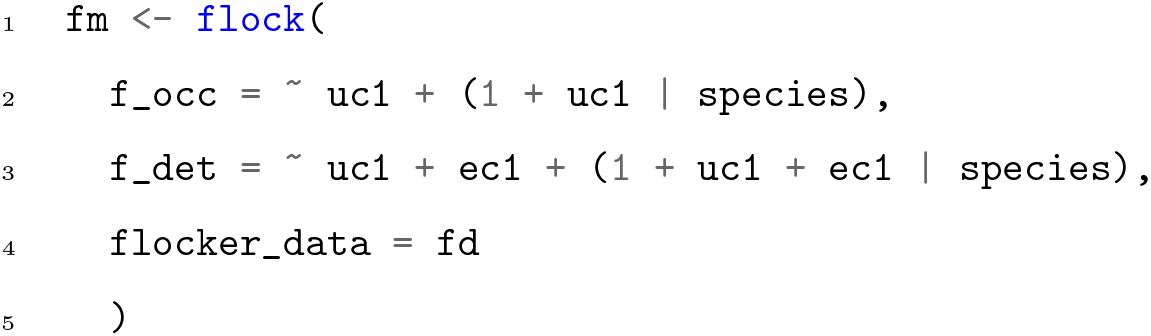

Arguments supplied to `flock()` define formulas using `brms` syntax for the occupancy (`f_occ`) and detection (`f_det`) components, and also provide the formatted data. At this stage, the full flexibility and power of `brms` formula syntax are available to the user.

## 3 Available models in flocker

### 3.1 The single-season rep-varying model

The above example shows the single-season rep-varying model (Box 1). The likelihood *ℒ* for a single closure-unit *i* is

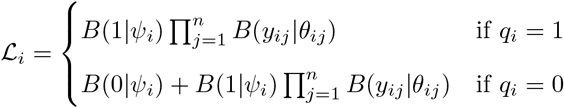

where *i* indexes the closure-unit and *j* indexes the replicate. *y*_*ij*_ is the observed data, *B*(*a*|*b*) is the Bernoulli probability mass function for datum *a* given probability *b, ψ*_*i*_ is the fitted occupancy probability at the closure-unit, *θ*_*ij*_ is the fitted detection probability for the visit, and *q*_*i*_ is an indicator that takes the value 1 if there is at least one detection at the closure-unit and 0 otherwise.

### 3.2 The single-season rep-constant model

In a single-season rep-constant model (i.e. one without event covariates; Box 1), a zero-inflated binomial likelihood provides a sufficient statistic that factorizes at the row level. This formulation improves efficiency and enables `brms` features including within-chain parallelization of model fitting and several post-processing functions for the output (see Post-processing below). Though `brms` comes with a native zero-inflated binomial distribution, `flocker` re-implements this distribution so that the linear predictor for occupancy represents an occupancy probability rather than a non-occupancy probability.

The likelihood is:

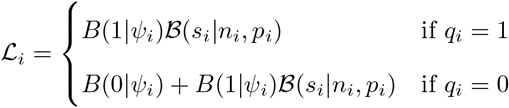

where *B*(*d*|*e, f*) is the binomial probability mass function for number of successes *d* given number of trials *e* and probability of success *f, s*_*i*_ is the number of visits with detections, *n*_*i*_ the total number of visits, *p*_*i*_ is the fitted detection probability applied to all repeat visits in the closure-unit. All other notation is carried over from the rep-varying model above.

### 3.3 Multi-season models

Multi-season occupancy models group closure-units within *series* that evolve via modeled colonization and extinction processes (*15*). `flocker` implements multi-season models using a hidden Markov model (HMM) approach (*18* –*20*). Note that efficient marginalized formulations of HMM likelihoods need not sacrifice the ability to perform inference on the unobserved occupancy states, *contra* (*21*).

The HMM conceptualizes the latent occupancy dynamics as a Markov-like process (optionally with time-varying transition probabilities (*20*)), where the transition probabilities between the two possible states (occupied or unoccupied) are a colonization probability *𝒞*, an extinction probability *ε*, and their complements (*19, 21*). The HMM involves a model for *𝒞*, a model for *ε*, and a model for the initial occupancy probability *𝒪* in the first timestep. We also need a model for the probability of observing the data in any particular timestep conditional on the true occupancy state in that timestep (in the HMM literature, this probability is referred to as an *emission probability*). These emission probabilities are identical to the detection sub-model in a single-season model.

`flocker` provides two approaches to modeling the colonization and extinction probabilities *𝒞* and *ε* and two approaches for modeling the initial occupancy probability *𝒪*, resulting in four flavors of multi-season model.

#### 3.3.1 Colonization and extinction

In *colonization-extinction* models, *𝒞* and *ε* are modeled directly (*15*). In *autologistic* models, the modeled occupancy probability in timestep *t* + 1 includes a term that depends on the latent true occupancy state at time *t* (*16, 22*). This autologistic term is generally modeled as zero when the site is unoccupied at time *t*, and some fitted constant when the site is occupied at time *t*. When the autologistic term is zero, the linear predictor gives a colonization probability (the probability of occupancy conditional on non-occupancy during the previous timestep). When the autologistic term is nonzero, the linear predictor gives a persistence probability *𝒫* = 1 *− ε*.

Thus, the autologistic model differs from the colonization-extinction model in that it assumes that *𝒞* and *ε* are related to one another via an offset that relates colonization to persistence. If the offset is modeled as constant, then this model ascribes a “suitability” to each site that defines the position of a pair of probabilities *𝒞* and *𝒫* that differ by a constant on the logit scale. `flocker` also provides the option to model the offset via its own covariate-based predictor, which reintroduces flexibility in the relationship between *𝒫* and *𝒞*, but tends to retain the idea that these probabilities are related and move in tandem.

#### 3.3.2 Initial occupancy

Most multi-season occupancy models estimate initial occupancy probabilities *𝒪* explicitly via their own predictor (*15, 16, 22*). Borrowing from the HMM literature, `flocker` provides optional functionality to handle *𝒪* differently (*20*). Any pair of time-invariant colonization and extinction probabilities defines an equilibrium proportion of occupied sites. If *𝒞* and *ε* remain constant for a long time prior to the first timestep, the initial occupancy probability should be this equilibrium frequency, eliminating the need to parameterize and fit *𝒪* separately from *𝒞* and *ε*.

This approach is sometimes valid even if *𝒞* and *ε* are time-varying, as long as they are invariant for a substantial period leading up to the first modeled timestep (and subsequent variation is modeled adequately). For this reason, `flocker` always computes the equilibrium based on the *𝒞* and *ε* modeled for the first timestep.

`flocker` does not provide an option to enforce equality between initial occupancy probabilities *𝒪* and colonization probabilities *𝒞* (*23, 24*). Although the autologistic model has the structural form of a single-season occupancy model plus an autologistic term, the colonization probability estimated by fixing the autologistic term to zero is not an estimate of the single-season occupancy probability. For data too scant to independently estimate *𝒪, 𝒞*, and an autologistic term, we recommend using the equilibrium initialization discussed above, which is as parsimonious as enforcing equality between *𝒪* and *𝒞* but reflects much more palatable assumptions.

#### 3.3.3 The generalized multi-season likelihood

For a multi-season model, we marginalize over the possible sequences of latent states within each series using the forward algorithm (*20*). The likelihood *ℳ* for a single series *m* of dynamically linked closure-units is

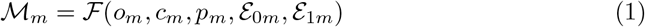

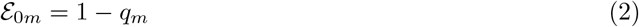

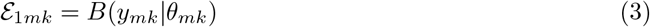

where *ℱ* is the forward algorithm, *o*_*m*_ is the initial occupancy probability, *c*_*m*_ is a vector of colonization probabilities, *p*_*m*_ is a vector of persistence probabilities, *ε*_0*m*_ is a vector of emission likelihoods conditional on non-occupancy, *ε*_1*m*_ is a vector of emission likelihoods conditional on occupancy, *q*_*m*_ is a vector of *q*_*i*_ for *i* corresponding to timeseries *m, y*_*mk*_ and *θ*_*mk*_ are vectors of *y* and *θ* for unit/visit indices pertaining to the *k*th timestep within timeseries *m*, and all other notation is carried over from the single-season models above.

### 3.4 Data-augmented multi-species models

Multi-species models can be extended via data augmentation to estimate occupancy of never-detected species (*17*). A multi-species dataset is augmented with a large number of never-detected pseudospecies, a parameter Ω gives the Bernoulli probability that any given (pseudo)species occurs in the study area, and random-effects exchangeability assumptions are applied to any species-specific parameters.

We marginalize over the occupancy status of a closure-unit as for a single-season model, yielding the unit-wise likelihood *ℒ*_*i*_. We additionally marginalize over the availability of each (pseudo)species, yielding the species-wise likelihood

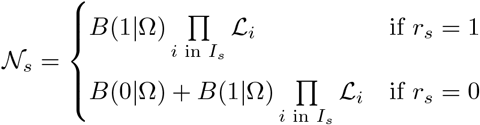

where *s* indexes the species, Ω is the fitted availability probability, *I*_*s*_ is the set of all closure-unit indices *i* pertaining to species *s, r*_*s*_ is an indicator that takes the value 1 if there is at least one positive detection of the (pseudo)species in the entire dataset and 0 otherwise, and all other notation is carried over from the models above.

`flocker` enables users to fit data-augmented models using arbitrary `brms` formulas for occupancy and detection. However, continuous covariates in the occupancy sub-model can lead to pitfalls in interpretation, because the model extrapolates about never-observed species that occur beyond the range of covariate values actually sampled (*25*). This extrapolation is unprincipled, most obviously when it fails to account for physical limits of a gradient (e.g. valley floor and mountain peak on an elevational gradient (*26*)). Following (*27*), we recommend extreme caution when interpreting patterns estimated for never-observed species.

## 4 Post-processing

Some `brms` post-processing functions, such as `summary()`, work for all `flocker` models. Other functionality works only for the rep-constant single-season model (which factorizes at the row level). `flocker` re-implements essential post-processing functionality for other model types; these functions also work with the rep-constant single-season model.

### 4.1 Fitted values

`fitted_flocker()` computes fitted values for the linear predictors, returning in the shape of the observation data inputted to `make_flocker_data()` . Although `brms::posterior_linpred()` runs on all `flocker` models, it returns in an obscure shape due to the vicissitudes of the `flocker` data format.

### 4.2 The posterior latent occupancy state

`get_Z()` extracts the posterior occupancy state from a `flockerfit` object. By default, `get_Z()` computes the posterior distribution conditional on the observed detection histories (closure-units with detections are occupied with certainty; closure-units without detections are occupied with probability less than the fitted occupancy probability). In multi-season models, the posterior conditions jointly on all observations within a series, via the forward-backward algorithm (*20*). In data-augmented models, the posterior conditions on whether a (pseudo)species has been detected at least once. Optionally, via the argument `history_condition = FALSE`, `get_Z()` can compute the posterior while conditioning on the fitted parameters but not on the observed detection histories.

By default `get_Z()` returns marginal occupancy probabilities, but by passing the argument `sample = TRUE` it will return Bernoulli samples from these probabilities. This is nontrivial for multi-season models, where we wish to return valid sequences of samples from the joint posterior. When `history_condition = FALSE`, we sample the occupancy state on a forward pass. When `history_condition = TRUE`, we sample the states on a backward pass (i.e. forward-filtering-backward-sampling rather than forward-filtering-backward-smoothing).

### 4.3 Posterior prediction

`predict_flocker()` replaces `brms::predict.brmsfit()` for posterior prediction of visit-level detection/non-detection data from a `flockerfit` object. Under the hood, predictions involve calling `get_Z(…, sample = TRUE)`, and the same considerations about `history_condition` apply. Importantly `history_condition` defaults to `FALSE` in posterior prediction, a *different default* from `get_Z()` . The reason for these different defaults is because `get_Z()` is commonly used for inference about the occupancy state at the study sites, whereas `predict_flocker()` is commonly used for posterior predictive checking. It is anti-conservative and undesirable to equip posterior predictive checks with perfect information about whether each closure-unit has at least one detection.

### 4.4 Cross-validation

`loo_compare_flocker()` provides functionality for model comparison based on computationally efficient approximate leave-one-out cross-validation via Pareto-smoothed importance sampling (PSIS-LOO) from R package `loo` (*28*). For all classes of model, we use withholds at the smallest level where the likelihood factorizes, thereby avoiding conceptual and computational challenges that bedevil all but a few non-factorizable models (*29*). Thus, for single-season models we perform leave-one-unit-out cross-validation. For multi-season models, we perform leave-one-series-out cross-validation. For multi-species models with data augmentation, we perform leave-one-(pseudo)species-out cross-validation.

## 5 Conclusions and future directions

`flocker` delivers the full flexibility of `brms` formula syntax to occupancy models and provides functionality for data simulation, data formatting, model fitting, posterior prediction, and model comparison. In the course of developing this package, we developed a vocabulary (*closure-unit, history-conditioning, unit covariate, event covariate*) that enables generalized discussion of the multiple models under the occupancy model umbrella. We hope that our unified treatment of these models and our user-friendly implementation will clarify and facilitate best practices.

One of the most exciting avenues for future improvements to `flocker` is the ongoing development of Stan and `brms`, which `flocker` will inherit. Further avenues for development on `flocker` include functionality for posterior predictive checking, within-chain parallelization for all model types, and additional likelihood types. We welcome issues and contributions at github.com/jsocolar/flocker.

## Supporting information

Supplemental SBC results

## 6 Authors’ contributions

JBS and SCM developed the package and wrote the manuscript.

## 7 Acknowledgments

We thank Paul Bürkner for insight into how to effectively harness `brms` for occupancy modeling via non-looped custom families. JBS thanks NCX for the space to pursue open-source development on `flocker` during his employment there.

