## Supplemental SBC results for "Introducing flocker: an R package for flexible occupancy modeling via brms and Stan": sbc.html

SBC for flocker models


### SBC for flocker models

###### Jacob Socolar

#### 2023-10-22

```
params <- list(
  n_sims = 1000,
  n_sites = 200,
  n_sims_augmented = 200,
  n_sites_augmented = 50,
  n_pseudospecies_augmented = 50
)
```

```
library(flocker)
library(brms)
library(SBC)
set.seed(1)
```

#### Overview

This document performs simulation-based calibration for the models
available in R package `flocker`. Here, our goal is to
validate `flocker`’s data formatting, decoding, and
likelihood implementations, and not `brms`’s construction of
the linear predictors.

The encoding of the data for a `flocker` model tends to be
more complex in the presence of missing observations, and so we include
missingness in the data simulation wherever possible (some visits
missing in all models, some time-steps missing in multiseason
models).

In all models, we include one unit covariate that affects detection
and occupancy, colonization, extinction and/or autologistic terms as
applicable, and one event covariate that affects detection only (for all
models except the rep-constant).

#### Single-season

##### Rep-constant

```
# make the stancode
model_name <- paste0(tempdir(), "/sbc_rep_constant_model.stan")
fd <- simulate_flocker_data(
  n_pt = params$n_sites, n_sp = 1,
  params = list(
    coefs = data.frame(
      det_intercept = rnorm(1),
      det_slope_unit = rnorm(1),
      occ_intercept = rnorm(1),
      occ_slope_unit = rnorm(1)
    )
  ),
  seed = NULL,
  rep_constant = TRUE,
  ragged_rep = TRUE
)
flocker_data = make_flocker_data(fd$obs, fd$unit_covs, quiet = TRUE)
  
scode <- flocker_stancode(
    f_occ = ~ 0 + Intercept + uc1,
    f_det = ~ 0 + Intercept + uc1,
    flocker_data = flocker_data,
    prior = 
      brms::set_prior("std_normal()") + 
      brms::set_prior("std_normal()", dpar = "occ"),
    backend = "cmdstanr"
  )

writeLines(scode, model_name)

rep_constant_generator <- function(N){  
  fd <- simulate_flocker_data(
    n_pt = N, n_sp = 1,
    params = list(
      coefs = data.frame(
        det_intercept = rnorm(1),
        det_slope_unit = rnorm(1),
        occ_intercept = rnorm(1),
        occ_slope_unit = rnorm(1)
      )
    ),
    seed = NULL,
    rep_constant = TRUE,
    ragged_rep = TRUE
  )
  
  flocker_data = make_flocker_data(fd$obs, fd$unit_covs, quiet = TRUE)
  
  # format for return
  list(
    variables = list(
      `b[1]` = fd$params$coefs$det_intercept,
      `b[2]` = fd$params$coefs$det_slope_unit,
      `b_occ[1]` = fd$params$coefs$occ_intercept,
      `b_occ[2]` = fd$params$coefs$occ_slope_unit
    ),
    generated = flocker_standata(
      f_occ = ~ 0 + Intercept + uc1,
      f_det = ~ 0 + Intercept + uc1,
      flocker_data = flocker_data
    )
  )
}

rep_constant_gen <- SBC_generator_function(
  rep_constant_generator, 
  N = params$n_sites
  )
rep_constant_dataset <- suppressMessages(
  generate_datasets(rep_constant_gen, params$n_sims)
)
  
rep_constant_backend <- 
  SBC_backend_cmdstan_sample(
    cmdstanr::cmdstan_model(
      paste0(tempdir(), "/sbc_rep_constant_model.stan")
      )
    )

rep_constant_results <- compute_SBC(rep_constant_dataset, rep_constant_backend)

plot_ecdf(rep_constant_results)
```

plot of chunk rep-constant

```
plot_rank_hist(rep_constant_results)
```

plot of chunk rep-constant

```
plot_ecdf_diff(rep_constant_results)
```

plot of chunk rep-constant

##### Rep-varying

```
# make the stancode
model_name <- paste0(tempdir(), "/sbc_rep_varying_model.stan")
fd <- simulate_flocker_data(
  n_pt = params$n_sites, n_sp = 1,
  params = list(
    coefs = data.frame(
      det_intercept = rnorm(1),
      det_slope_unit = rnorm(1),
      det_slope_visit = rnorm(1),
      occ_intercept = rnorm(1),
      occ_slope_unit = rnorm(1)
    )
  ),
  seed = NULL,
  rep_constant = FALSE,
  ragged_rep = TRUE
)
flocker_data = make_flocker_data(fd$obs, fd$unit_covs, fd$event_covs, quiet = TRUE)
  
scode <- flocker_stancode(
    f_occ = ~ 0 + Intercept + uc1,
    f_det = ~ 0 + Intercept + uc1 + ec1,
    flocker_data = flocker_data,
    prior = 
      brms::set_prior("std_normal()") + 
      brms::set_prior("std_normal()", dpar = "occ"),
    backend = "cmdstanr"
  )
writeLines(scode, model_name)

rep_varying_generator <- function(N){  
  fd <- simulate_flocker_data(
    n_pt = N, n_sp = 1,
    params = list(
      coefs = data.frame(
        det_intercept = rnorm(1),
        det_slope_unit = rnorm(1),
        det_slope_visit = rnorm(1),
        occ_intercept = rnorm(1),
        occ_slope_unit = rnorm(1)
      )
    ),
    seed = NULL,
    rep_constant = FALSE,
    ragged_rep = TRUE
  )
  
  flocker_data = make_flocker_data(fd$obs, fd$unit_covs, fd$event_covs, quiet = TRUE)
  
  # format for return
  list(
    variables = list(
      `b[1]` = fd$params$coefs$det_intercept,
      `b[2]` = fd$params$coefs$det_slope_unit,
      `b[3]` = fd$params$coefs$det_slope_visit,
      `b_occ[1]` = fd$params$coefs$occ_intercept,
      `b_occ[2]` = fd$params$coefs$occ_slope_unit
    ),
    generated = flocker_standata(
      f_occ = ~ 0 + Intercept + uc1,
      f_det = ~ 0 + Intercept + uc1 + ec1,
      flocker_data = flocker_data
    )
  )
}

rep_varying_gen <- SBC_generator_function(
  rep_varying_generator, 
  N = params$n_sites
  )
rep_varying_dataset <- suppressMessages(
  generate_datasets(rep_varying_gen, params$n_sims)
)
  
rep_varying_backend <- 
  SBC_backend_cmdstan_sample(
    cmdstanr::cmdstan_model(
      paste0(tempdir(), "/sbc_rep_varying_model.stan")
      )
    )

rep_varying_results <- compute_SBC(rep_varying_dataset, rep_varying_backend)
```

```
##  - 3 (0%) fits had at least one Rhat > 1.01. Largest Rhat was 1.016.
```

```
## Not all diagnostics are OK.
## You can learn more by inspecting $default_diagnostics, $backend_diagnostics 
## and/or investigating $outputs/$messages/$warnings for detailed output from the backend.
```

```
plot_ecdf(rep_varying_results)
```

plot of chunk rep-varying

```
plot_rank_hist(rep_varying_results)
```

plot of chunk rep-varying

```
plot_ecdf_diff(rep_varying_results)
```

plot of chunk rep-varying

#### Multi-season

`flocker` fits multi-season models that parameterize the
dynamics using colonization/extinction or autologistic specifications,
and that parameterize the initial occupancy state using explicit and
equilibrium parameterizations, for a total of four classes of
multi-season model. We validate each class.

##### Colonization-extinction, explicit initial occupancy

```
# make the stancode
model_name <- paste0(tempdir(), "/sbc_colex_ex_model.stan")
fd <- simulate_flocker_data(
  n_pt = params$n_sites, n_sp = 1, n_season = 4,
  params = list(
    coefs = data.frame(
      det_intercept = rnorm(1),
      det_slope_unit = rnorm(1),
      det_slope_visit = rnorm(1),
      occ_intercept = rnorm(1),
      occ_slope_unit = rnorm(1),
      col_intercept = rnorm(1),
      col_slope_unit = rnorm(1),
      ex_intercept = rnorm(1),
      ex_slope_unit = rnorm(1)
    )
  ),
  seed = NULL,
  rep_constant = FALSE,
  multiseason = "colex",
  multi_init = "explicit",
  ragged_rep = TRUE
)
flocker_data = make_flocker_data(
  fd$obs, fd$unit_covs, fd$event_covs,
  type = "multi", quiet = TRUE)
  
scode <- flocker_stancode(
    f_occ = ~ 0 + Intercept + uc1,
    f_col = ~ 0 + Intercept + uc1,
    f_ex = ~ 0 + Intercept + uc1,
    f_det = ~ 0 + Intercept + uc1 + ec1,
    flocker_data = flocker_data,
    prior = 
      brms::set_prior("std_normal()") + 
      brms::set_prior("std_normal()", dpar = "occ") +
      brms::set_prior("std_normal()", dpar = "colo") +
      brms::set_prior("std_normal()", dpar = "ex"),
    multiseason = "colex",
    multi_init = "explicit",
    backend = "cmdstanr"
  )
writeLines(scode, model_name)

colex_ex_generator <- function(N){  
  fd <- simulate_flocker_data(
    n_pt = params$n_sites, n_sp = 1, n_season = 4,
    params = list(
        det_intercept = rnorm(1),
        det_slope_unit = rnorm(1),
        det_slope_visit = rnorm(1),
        occ_intercept = rnorm(1),
        occ_slope_unit = rnorm(1),
        colo_intercept = rnorm(1),
        colo_slope_unit = rnorm(1),
        ex_intercept = rnorm(1),
        ex_slope_unit = rnorm(1)
    ),
    seed = NULL,
    rep_constant = FALSE,
    multiseason = "colex",
    multi_init = "explicit",
    ragged_rep = TRUE
  )
  
  flocker_data = make_flocker_data(
    fd$obs, fd$unit_covs, fd$event_covs,
    type = "multi", quiet = TRUE)
  
  # format for return
  list(
    variables = list(
      `b[1]` = fd$params$coefs$det_intercept,
      `b[2]` = fd$params$coefs$det_slope_unit,
      `b[3]` = fd$params$coefs$det_slope_visit,
      `b_occ[1]` = fd$params$coefs$occ_intercept,
      `b_occ[2]` = fd$params$coefs$occ_slope_unit,
      `b_colo[1]` = fd$params$coefs$col_intercept,
      `b_colo[2]` = fd$params$coefs$col_slope_unit,
      `b_ex[1]` = fd$params$coefs$ex_intercept,
      `b_ex[2]` = fd$params$coefs$ex_slope_unit
    ),
    generated = flocker_standata(
      f_occ = ~ 0 + Intercept + uc1,
      f_col = ~ 0 + Intercept + uc1,
      f_ex = ~ 0 + Intercept + uc1,
      f_det = ~ 0 + Intercept + uc1 + ec1,
      flocker_data = flocker_data,
      multiseason = "colex",
      multi_init = "explicit"
    )
  )
}

colex_ex_gen <- SBC_generator_function(
  colex_ex_generator, 
  N = params$n_sites
  )
colex_ex_dataset <- suppressMessages(
  generate_datasets(colex_ex_gen, params$n_sims)
)
  
colex_ex_backend <- 
  SBC_backend_cmdstan_sample(
    cmdstanr::cmdstan_model(
      paste0(tempdir(), "/sbc_colex_ex_model.stan")
      )
    )

colex_ex_results <- compute_SBC(colex_ex_dataset, colex_ex_backend)

plot_ecdf(colex_ex_results)
```

plot of chunk multi-colex-ex

```
plot_rank_hist(colex_ex_results)
```

plot of chunk multi-colex-ex

```
plot_ecdf_diff(colex_ex_results)
```

plot of chunk multi-colex-ex

##### Colonization-extinction, equilibrium initial occupancy

```
# make the stancode
model_name <- paste0(tempdir(), "/sbc_colex_eq_model.stan")
fd <- simulate_flocker_data(
  n_pt = params$n_sites, n_sp = 1, n_season = 4,
  params = list(
      det_intercept = rnorm(1),
      det_slope_unit = rnorm(1),
      det_slope_visit = rnorm(1),
      colo_intercept = rnorm(1),
      colo_slope_unit = rnorm(1),
      ex_intercept = rnorm(1),
      ex_slope_unit = rnorm(1)
  ),
  seed = NULL,
  rep_constant = FALSE,
  multiseason = "colex",
  multi_init = "equilibrium",
  ragged_rep = TRUE
)
flocker_data = make_flocker_data(
  fd$obs, fd$unit_covs, fd$event_covs,
  type = "multi", quiet = TRUE)
  
scode <- flocker_stancode(
  f_col = ~ 0 + Intercept + uc1,
  f_ex = ~ 0 + Intercept + uc1,
  f_det = ~ 0 + Intercept + uc1 + ec1,
  flocker_data = flocker_data,
  prior = 
    brms::set_prior("std_normal()") + 
    brms::set_prior("std_normal()", dpar = "colo") +
    brms::set_prior("std_normal()", dpar = "ex"),
  multiseason = "colex",
  multi_init = "equilibrium",
  backend = "cmdstanr"
  )
writeLines(scode, model_name)

colex_eq_generator <- function(N){  
  fd <- simulate_flocker_data(
    n_pt = params$n_sites, n_sp = 1, n_season = 4,
    params = list(
        det_intercept = rnorm(1),
        det_slope_unit = rnorm(1),
        det_slope_visit = rnorm(1),
        col_intercept = rnorm(1),
        col_slope_unit = rnorm(1),
        ex_intercept = rnorm(1),
        ex_slope_unit = rnorm(1)
    ),
    seed = NULL,
    rep_constant = FALSE,
    multiseason = "colex",
    multi_init = "equilibrium",
    ragged_rep = TRUE
  )
  
  flocker_data = make_flocker_data(
    fd$obs, fd$unit_covs, fd$event_covs,
    type = "multi", quiet = TRUE)
  
  # format for return
  list(
    variables = list(
      `b[1]` = fd$params$coefs$det_intercept,
      `b[2]` = fd$params$coefs$det_slope_unit,
      `b[3]` = fd$params$coefs$det_slope_visit,
      `b_colo[1]` = fd$params$coefs$col_intercept,
      `b_colo[2]` = fd$params$coefs$col_slope_unit,
      `b_ex[1]` = fd$params$coefs$ex_intercept,
      `b_ex[2]` = fd$params$coefs$ex_slope_unit
    ),
    generated = flocker_standata(
      f_col = ~ 0 + Intercept + uc1,
      f_ex = ~ 0 + Intercept + uc1,
      f_det = ~ 0 + Intercept + uc1 + ec1,
      flocker_data = flocker_data,
      multiseason = "colex",
      multi_init = "equilibrium"
    )
  )
}

colex_eq_gen <- SBC_generator_function(
  colex_eq_generator, 
  N = params$n_sites
  )
colex_eq_dataset <- suppressMessages(
  generate_datasets(colex_eq_gen, params$n_sims)
)
  
colex_eq_backend <- 
  SBC_backend_cmdstan_sample(
    cmdstanr::cmdstan_model(
      paste0(tempdir(), "/sbc_colex_eq_model.stan")
      )
    )

colex_eq_results <- compute_SBC(colex_eq_dataset, colex_eq_backend)
```

```
##  - 1 (0%) fits had at least one Rhat > 1.01. Largest Rhat was 1.012.
```

```
##  - 853 (85%) fits had some steps rejected. Maximum number of rejections was 5.
```

```
## Not all diagnostics are OK.
## You can learn more by inspecting $default_diagnostics, $backend_diagnostics 
## and/or investigating $outputs/$messages/$warnings for detailed output from the backend.
```

```
plot_ecdf(colex_eq_results)
```

plot of chunk multi-colex-eq

```
plot_rank_hist(colex_eq_results)
```

plot of chunk multi-colex-eq

```
plot_ecdf_diff(colex_eq_results)
```

plot of chunk multi-colex-eq

##### Autologistic, explicit initial occupancy

```
# make the stancode
model_name <- paste0(tempdir(), "/sbc_auto_ex_model.stan")
fd <- simulate_flocker_data(
  n_pt = params$n_sites, n_sp = 1, n_season = 4,
  params = list(
      det_intercept = rnorm(1),
      det_slope_unit = rnorm(1),
      det_slope_visit = rnorm(1),
      occ_intercept = rnorm(1),
      occ_slope_unit = rnorm(1),
      col_intercept = rnorm(1),
      col_slope_unit = rnorm(1),
      auto_intercept = rnorm(1),
      auto_slope_unit = rnorm(1)
  ),
  seed = NULL,
  rep_constant = FALSE,
  multiseason = "autologistic",
  multi_init = "explicit",
  ragged_rep = TRUE
)
flocker_data = make_flocker_data(
  fd$obs, fd$unit_covs, fd$event_covs,
  type = "multi", quiet = TRUE)
  
scode <- flocker_stancode(
    f_occ = ~ 0 + Intercept + uc1,
    f_col = ~ 0 + Intercept + uc1,
    f_auto = ~ 0 + Intercept + uc1,
    f_det = ~ 0 + Intercept + uc1 + ec1,
    flocker_data = flocker_data,
    prior = 
      brms::set_prior("std_normal()") + 
      brms::set_prior("std_normal()", dpar = "occ") +
      brms::set_prior("std_normal()", dpar = "colo") +
      brms::set_prior("std_normal()", dpar = "autologistic"),
    multiseason = "autologistic",
    multi_init = "explicit",
    backend = "cmdstanr"
  )
writeLines(scode, model_name)

auto_ex_generator <- function(N){  
  fd <- simulate_flocker_data(
    n_pt = params$n_sites, n_sp = 1, n_season = 4,
    params = list(
        det_intercept = rnorm(1),
        det_slope_unit = rnorm(1),
        det_slope_visit = rnorm(1),
        occ_intercept = rnorm(1),
        occ_slope_unit = rnorm(1),
        colo_intercept = rnorm(1),
        colo_slope_unit = rnorm(1),
        auto_intercept = rnorm(1),
        auto_slope_unit = rnorm(1)
    ),
    seed = NULL,
    rep_constant = FALSE,
    multiseason = "autologistic",
    multi_init = "explicit",
    ragged_rep = TRUE
  )
  
  flocker_data = make_flocker_data(
    fd$obs, fd$unit_covs, fd$event_covs,
    type = "multi", quiet = TRUE)
  
  # format for return
  list(
    variables = list(
      `b[1]` = fd$params$coefs$det_intercept,
      `b[2]` = fd$params$coefs$det_slope_unit,
      `b[3]` = fd$params$coefs$det_slope_visit,
      `b_occ[1]` = fd$params$coefs$occ_intercept,
      `b_occ[2]` = fd$params$coefs$occ_slope_unit,
      `b_colo[1]` = fd$params$coefs$col_intercept,
      `b_colo[2]` = fd$params$coefs$col_slope_unit,
      `b_autologistic[1]` = fd$params$coefs$auto_intercept,
      `b_autologistic[2]` = fd$params$coefs$auto_slope_unit
    ),
    generated = flocker_standata(
      f_occ = ~ 0 + Intercept + uc1,
      f_col = ~ 0 + Intercept + uc1,
      f_auto = ~ 0 + Intercept + uc1,
      f_det = ~ 0 + Intercept + uc1 + ec1,
      flocker_data = flocker_data,
      multiseason = "autologistic",
      multi_init = "explicit"
    )
  )
}

auto_ex_gen <- SBC_generator_function(
  auto_ex_generator, 
  N = params$n_sites
  )
auto_ex_dataset <- suppressMessages(
  generate_datasets(auto_ex_gen, params$n_sims)
)
  
auto_ex_backend <- 
  SBC_backend_cmdstan_sample(
    cmdstanr::cmdstan_model(
      paste0(tempdir(), "/sbc_auto_ex_model.stan")
      )
    )

auto_ex_results <- compute_SBC(auto_ex_dataset, auto_ex_backend)

plot_ecdf(auto_ex_results)
```

plot of chunk multi-auto-ex

```
plot_rank_hist(auto_ex_results)
```

plot of chunk multi-auto-ex

```
plot_ecdf_diff(auto_ex_results)
```

plot of chunk multi-auto-ex

##### Autologistic, equilibrium initial occupancy

```
# make the stancode
model_name <- paste0(tempdir(), "/sbc_auto_eq_model.stan")
fd <- simulate_flocker_data(
  n_pt = params$n_sites, n_sp = 1, n_season = 4,
  params = list(
      det_intercept = rnorm(1),
      det_slope_unit = rnorm(1),
      det_slope_visit = rnorm(1),
      auto_intercept = rnorm(1),
      auto_slope_unit = rnorm(1)
  ),
  seed = NULL,
  rep_constant = FALSE,
  multiseason = "autologistic",
  multi_init = "equilibrium",
  ragged_rep = TRUE
)
flocker_data = make_flocker_data(
  fd$obs, fd$unit_covs, fd$event_covs,
  type = "multi", quiet = TRUE)
  
scode <- flocker_stancode(
    f_col = ~ 0 + Intercept + uc1,
    f_auto = ~ 0 + Intercept + uc1,
    f_det = ~ 0 + Intercept + uc1 + ec1,
    flocker_data = flocker_data,
    prior = 
      brms::set_prior("std_normal()") + 
      brms::set_prior("std_normal()", dpar = "colo") +
      brms::set_prior("std_normal()", dpar = "autologistic"),
    multiseason = "autologistic",
    multi_init = "equilibrium",
    backend = "cmdstanr"
  )
writeLines(scode, model_name)

auto_eq_generator <- function(N){  
  fd <- simulate_flocker_data(
    n_pt = params$n_sites, n_sp = 1, n_season = 4,
    params = list(
        det_intercept = rnorm(1),
        det_slope_unit = rnorm(1),
        det_slope_visit = rnorm(1),
        col_intercept = rnorm(1),
        col_slope_unit = rnorm(1),
        auto_intercept = rnorm(1),
        auto_slope_unit = rnorm(1)
    ),
    seed = NULL,
    rep_constant = FALSE,
    multiseason = "autologistic",
    multi_init = "equilibrium",
    ragged_rep = TRUE
  )
  
  flocker_data = make_flocker_data(
    fd$obs, fd$unit_covs, fd$event_covs,
    type = "multi", quiet = TRUE)
  
  # format for return
  list(
    variables = list(
      `b[1]` = fd$params$coefs$det_intercept,
      `b[2]` = fd$params$coefs$det_slope_unit,
      `b[3]` = fd$params$coefs$det_slope_visit,
      `b_colo[1]` = fd$params$coefs$col_intercept,
      `b_colo[2]` = fd$params$coefs$col_slope_unit,
      `b_autologistic[1]` = fd$params$coefs$auto_intercept,
      `b_autologistic[2]` = fd$params$coefs$auto_slope_unit
    ),
    generated = flocker_standata(
      f_col = ~ 0 + Intercept + uc1,
      f_auto = ~ 0 + Intercept + uc1,
      f_det = ~ 0 + Intercept + uc1 + ec1,
      flocker_data = flocker_data,
      multiseason = "autologistic",
      multi_init = "equilibrium"
    )
  )
}

auto_eq_gen <- SBC_generator_function(
  auto_eq_generator, 
  N = params$n_sites
  )
auto_eq_dataset <- suppressMessages(
  generate_datasets(auto_eq_gen, params$n_sims)
)
  
auto_eq_backend <- 
  SBC_backend_cmdstan_sample(
    cmdstanr::cmdstan_model(
      paste0(tempdir(), "/sbc_auto_eq_model.stan")
      )
    )

auto_eq_results <- compute_SBC(auto_eq_dataset, auto_eq_backend)
```

```
##  - 302 (30%) fits had some steps rejected. Maximum number of rejections was 3.
```

```
## Not all diagnostics are OK.
## You can learn more by inspecting $default_diagnostics, $backend_diagnostics 
## and/or investigating $outputs/$messages/$warnings for detailed output from the backend.
```

```
plot_ecdf(auto_eq_results)
```

plot of chunk multi-auto-eq

```
plot_rank_hist(auto_eq_results)
```

plot of chunk multi-auto-eq

```
plot_ecdf_diff(auto_eq_results)
```

plot of chunk multi-auto-eq

#### Data-augmented

```
# make the stancode
model_name <- paste0(tempdir(), "/sbc_augmented_model.stan")

omega <- boot::inv.logit(rnorm(1, 0, .1))
available <- rbinom(1, params$n_pseudospecies_augmented, omega)
unavailable <- params$n_pseudospecies_augmented - available
coef_means <- rnorm(5) # normal prior on random effect means
sigma <- abs(rnorm(5)) # half-normal prior on all random effect sds
Sigma <- diag(5) * sigma

fd <- simulate_flocker_data(
  n_pt = params$n_sites_augmented, n_sp = available,
  params = list(
    coef_means = coef_means,
    Sigma = Sigma,
    coefs = data.frame(
      det_intercept = rnorm(available, coef_means[1], sigma[1]),
      det_slope_unit = rnorm(available, coef_means[2], sigma[2]),
      det_slope_visit = rnorm(available, coef_means[3], sigma[3]),
      occ_intercept = rnorm(available, coef_means[4], sigma[4]),
      occ_slope_unit = rnorm(available, coef_means[5], sigma[5])
    )
  ),
  seed = NULL,
  rep_constant = FALSE,
  ragged_rep = TRUE
)

obs_aug <- fd$obs[seq_len(params$n_sites_augmented), ]

for(i in 2:available){
  obs_aug <- abind::abind(
    obs_aug, 
    fd$obs[((i - 1) * params$n_sites_augmented) + seq_len(params$n_sites_augmented), ], 
    along = 3
    )
}

event_covs_aug <- list(ec1 = fd$event_covs$ec1[seq_len(params$n_sites_augmented), ])
unit_covs_aug <- data.frame(uc1 = fd$unit_covs[seq_len(params$n_sites_augmented), "uc1"])

flocker_data = make_flocker_data(
  obs_aug, unit_covs_aug, event_covs_aug,
  type = "augmented", n_aug = unavailable,
  quiet = TRUE)
  
scode <- flocker_stancode(
    f_occ = ~ 0 + Intercept + uc1 + (1 + uc1 || ff_species),
    f_det = ~ 0 + Intercept + uc1 + ec1 + (1 + uc1 + ec1 || ff_species),
    flocker_data = flocker_data,
    prior = 
      brms::set_prior("std_normal()") + 
      brms::set_prior("std_normal()", class = "sd") +
      brms::set_prior("std_normal()", dpar = "occ") +
      brms::set_prior("std_normal()", class = "sd", dpar = "occ") +
      brms::set_prior("normal(0, 0.1)", class = "Intercept", dpar = "Omega"),
    backend = "cmdstanr",
    augmented = TRUE
  )
writeLines(scode, model_name)

aug_generator <- function(N){  
  omega <- boot::inv.logit(rnorm(1, 0, .1))
  available <- rbinom(1, params$n_pseudospecies_augmented, omega)
  unavailable <- params$n_pseudospecies_augmented - available
  coef_means <- rnorm(5) # normal prior on random effect means
  sigma <- abs(rnorm(5)) # half-normal prior on all random effect sds
  Sigma <- diag(5) * sigma
  
  fd <- simulate_flocker_data(
    n_pt = params$n_sites_augmented, n_sp = available,
    params = list(
      coef_means = coef_means,
      Sigma = Sigma,
      coefs = data.frame(
        det_intercept = rnorm(available, coef_means[1], sigma[1]),
        det_slope_unit = rnorm(available, coef_means[2], sigma[2]),
        det_slope_visit = rnorm(available, coef_means[3], sigma[3]),
        occ_intercept = rnorm(available, coef_means[4], sigma[4]),
        occ_slope_unit = rnorm(available, coef_means[5], sigma[5])
      )
    ),
    seed = NULL,
    rep_constant = FALSE,
    ragged_rep = TRUE
  )
  
  obs_aug <- fd$obs[seq_len(params$n_sites_augmented), ]
  
  for(i in 2:available){
    obs_aug <- abind::abind(
      obs_aug, 
      fd$obs[((i - 1) * params$n_sites_augmented) + seq_len(params$n_sites_augmented), ], 
      along = 3
      )
  }
  
  event_covs_aug <- list(ec1 = fd$event_covs$ec1[seq_len(params$n_sites_augmented), ])
  unit_covs_aug <- data.frame(uc1 = fd$unit_covs[seq_len(params$n_sites_augmented), "uc1"])
  
  flocker_data = make_flocker_data(
    obs_aug, unit_covs_aug, event_covs_aug,
    type = "augmented", n_aug = unavailable,
    quiet = TRUE)
  # format for return
  list(
    variables = list(
      `b[1]` = fd$params$coef_means[1],
      `b[2]` = fd$params$coef_means[2],
      `b[3]` = fd$params$coef_means[3],
      `b_occ[1]` = fd$params$coef_means[4],
      `b_occ[2]` = fd$params$coef_means[5],
      `Intercept_Omega` = boot::logit(omega)
    ),
    generated = flocker_standata(
      f_occ = ~ 0 + Intercept + uc1 + (1 + uc1 || ff_species),
      f_det = ~ 0 + Intercept + uc1 + ec1 + (1 + uc1 + ec1 || ff_species),
      flocker_data = flocker_data,
      augmented = TRUE
    )
  )
}

aug_gen <- SBC_generator_function(
  aug_generator, 
  N = params$n_sites_augmented
  )
aug_dataset <- suppressMessages(
  generate_datasets(aug_gen, params$n_sims_augmented)
)
  
aug_backend <- 
  SBC_backend_cmdstan_sample(
    cmdstanr::cmdstan_model(
      paste0(tempdir(), "/sbc_augmented_model.stan")
      )
    )

aug_results <- compute_SBC(aug_dataset, aug_backend)
```

```
##  - 4 (2%) fits had at least one Rhat > 1.01. Largest Rhat was 1.022.
```

```
##  - 2 (1%) fits had divergent transitions. Maximum number of divergences was 2.
```

```
##  - 7 (4%) fits had some steps rejected. Maximum number of rejections was 1.
```

```
## Not all diagnostics are OK.
## You can learn more by inspecting $default_diagnostics, $backend_diagnostics 
## and/or investigating $outputs/$messages/$warnings for detailed output from the backend.
```

```
plot_ecdf(aug_results)
```

plot of chunk data-augmented

```
plot_rank_hist(aug_results)
```

plot of chunk data-augmented

```
plot_ecdf_diff(aug_results)
```

plot of chunk data-augmented
